# An Optimized Buffer for Repeatable Multicolor STORM

**DOI:** 10.1101/2022.05.19.491818

**Authors:** Vaky Abdelsayed, Hadjer Boukhatem, Nicolas Olivier

## Abstract

STORM microscopy is one of the most popular method of super-resolution microscopy, due to moderate requirements on the optical setup, and high achievable resolution. However, since its inception more than 15 years ago, protocols have barely evolved, and despite some recent progress, multicolor imaging can still be complex without the right equipment. We decided to optimize the buffer composition to improve the blinking of the most popular red dye CF-568 while maintaining good performance for far-red fluorophores such as Alexa-647 using the concentration of three chemicals and the pH as 4 optimization parameters. We developed a simple, cheap and stable buffer, that can be stored several weeks and frozen for longer term storage that allow high quality 3-color STORM imaging.

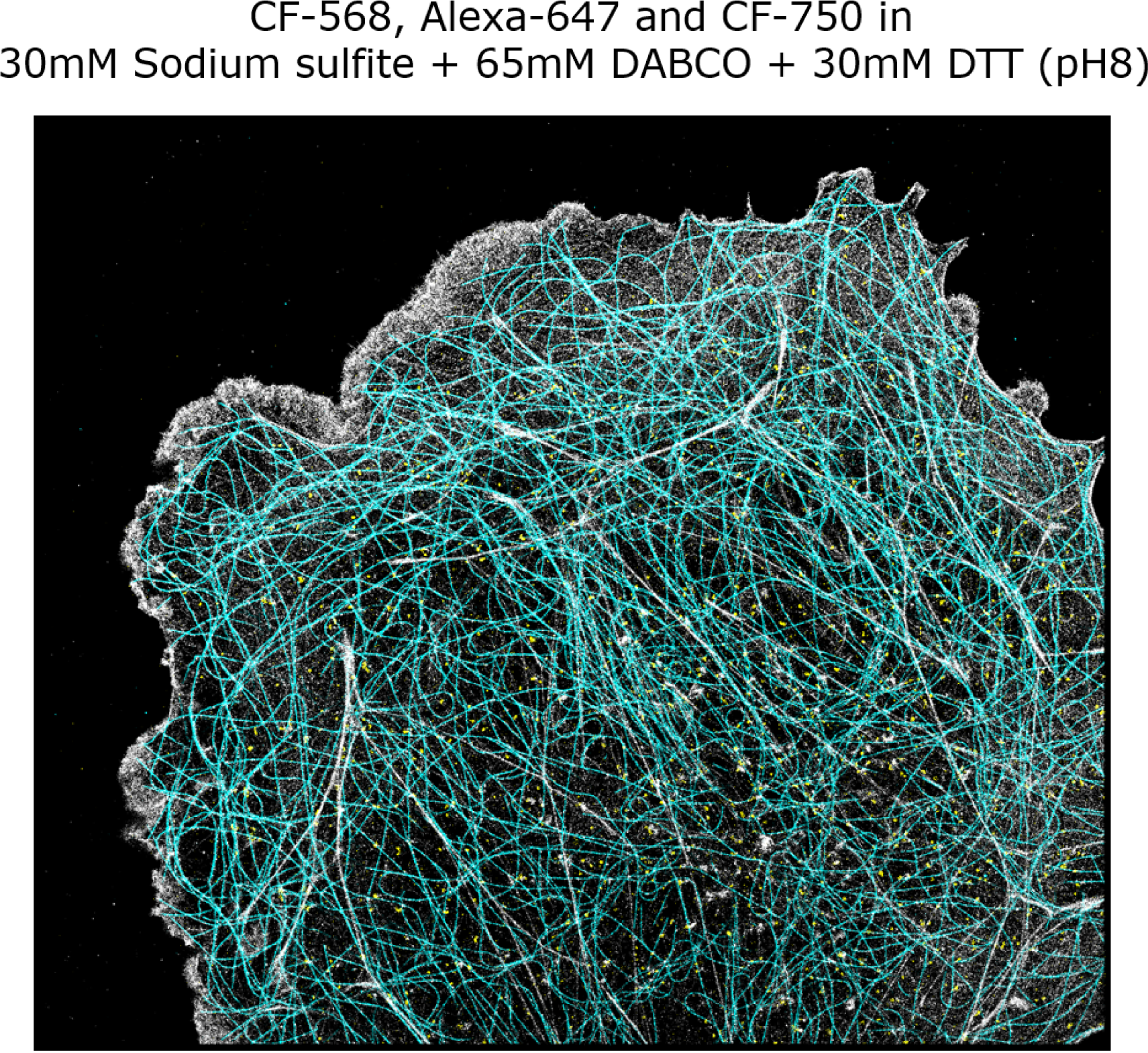

## Introduction

STORM microscopy (1) is a powerful optical super-resolution method, that can now yield sub-10 nm resolution in 3D for multiple colors on optimized optical setups (2). The key to this method is to induce reversible switching of (mainly organic) fluorophores by controlling their chemical environment (3–5). Most studies still rely on the original chemical environment: a combination of enzymatic oxygen scavenger (usually, Glucose-oxydase and catalase) and a reducing thiol such as 2-Mercaptoethylamine/Cysteamine (MEA) or *β*-Mercaptoethanol (*β*ME), as the blinking mechanism of the most popular fluorophores Cy5/Alexa-647 has been well-studied in this environment (6). Moreover, several screens have identified dyes that perform well in this buffer (7, 8), with the most promising 2-color combination being CF-568 and Alexa-647. Far-red fluorophores excited at 750 nm have also been used (7, 9) with success in combination with Alexa-647, but unfortunately most commercial microscopes do not offer this laser wavelength and lower photon counts for *≈*750-nm excited dyes than for Alexa-647 means the buffers have to be optimized specifically (9, 10). Another popular multicolor solution relies on simultaneous excitation of 640-nm excited dyes followed by spectral un-mixing (8, 11, 12), which can be used for up to 3 colors simultaneously, but require specific analysis software which only recently became publicly available (13, 14) and has limitations in terms of cross-talk for dense regions, and for the identification of structures of interest. In order to improve multicolor STORM imaging protocols, we decided to focus on CF-568, the most popular second color for STORM, and to optimize the buffers by quantifying the image quality of microtubules samples (see Methods). This approach does not provide photophysical parameters such as off-times, absolute numbers of blinking but does provide some quantitative measure of image quality. In terms of parameters to optimize for the buffer composition, we made the following decisions:

1. To use Sodium Sulfite as an oxygen scavenger (15). It is simple to use compared to enzymatic systems, reduces costs, and provides stability (both temporal, and in terms of pH).
2. To use DTT as the primary reducing agent. It can be purchased as a 1 M solution, so does not have to be prepared fresh from a powder as MEA, and is not as stinky nor as toxic as *β*ME.
3. To use DABCO (16) as an additional triplet-state quencher. DABCO was previously used to improve the blinking of Cy3 (17) and does not prevent the blinking of Alexa-647 (18).
4. To vary the pH, which is made simple by the lack of acidification, and the possibility to prepare large volumes of buffers to use a pH-meter.
5. To use water as the main solvent. While higher index media are very useful for 3D imaging (15, 17, 19) they make TIRF imaging more difficult, and the majority of STORM imaging is performed in TIRF or grazing incidence with TIRF-based z-stabilization.

We also decided to work on a fairly simple microscope (see Methods: epi-illumination using multimode lasers and fibers, multicolor dichroic and filters), and not to use any UV reactivation to make sure the protocols are generally applicable.

## Methods

### Optical Setup

Buffer optimization for CF-568 was performed on a home built microscope (see Fig. 1) based on the MiCube design (20), equipped with an Olympus 100x 1.45NA objective, and a 200 mm infinity-corrected tube lens (Thorlabs ITL200), resulting in 111*×* magnification on the camera (Andor iXon), for an effective pixel size of 144 nm. The sample was placed on a 300 *µ*m range z-piezo stage (LTZ-300, Piezoconcept) with manual lateral movement which is the only moving part of the system. We used a 5 color dichroic mirror and emission filter (Semrock FF409/493/573/652/759-Di01 + FF01-432/515/595/681/809), and added an additional filter to remove laser reflections (ET605/70m (Chroma) for CF-568). We used a 561 nm laser (100 mW, Cobolt) coupled to a 400 *µ*m Multi-Mode fiber (M28L02, Thorlabs) which was shaken using a fan to cancel out speckles, and the intensity at the sample from the 561 nm laser used for the buffer optimization was in the 1*−* 3*kW*.*cm*^*−*2^ range. Further imaging, including 2-color and 3-color STORM was performed on an IX83 Inverted microscope (Olympus) using a 100x 1.3NA objective (Olympus), and an Orca Fusion sCMOS camera (Hamamatsu) using the slowest read-out speed. We used a free-space 532 nm laser (Voltran, 40 mW), a 561 nm laser (100 mW, Cobolt), a 635 nm multimode laser (700 mW, Lasertak) and a 750 nm multimode (1.2 W Oxxius), the later 3 were coupled into a 400 *µ*m Multi-Mode fiber (M28L02, Thorlabs) which was shaken with a fan as above. The microscope is equipped with Chroma filters: 532 nm and 640 nm-excited fluorophores were imaged using a ZT532/640rpc 2-color dichroic mirror, and a ET605/70 or ET700-75 emission filter. Due to the double-deck design, the emitted light also goes through a T550lpxr dichroic or a T660lpxr dichroic. 750 nm-excited fluorophores were imaged using a T760lpxr dichroic, and a ET810-90 emission filter. Similarly, the emitted light also goes through an additional T760lpxr dichroic.

**Fig. 1.**
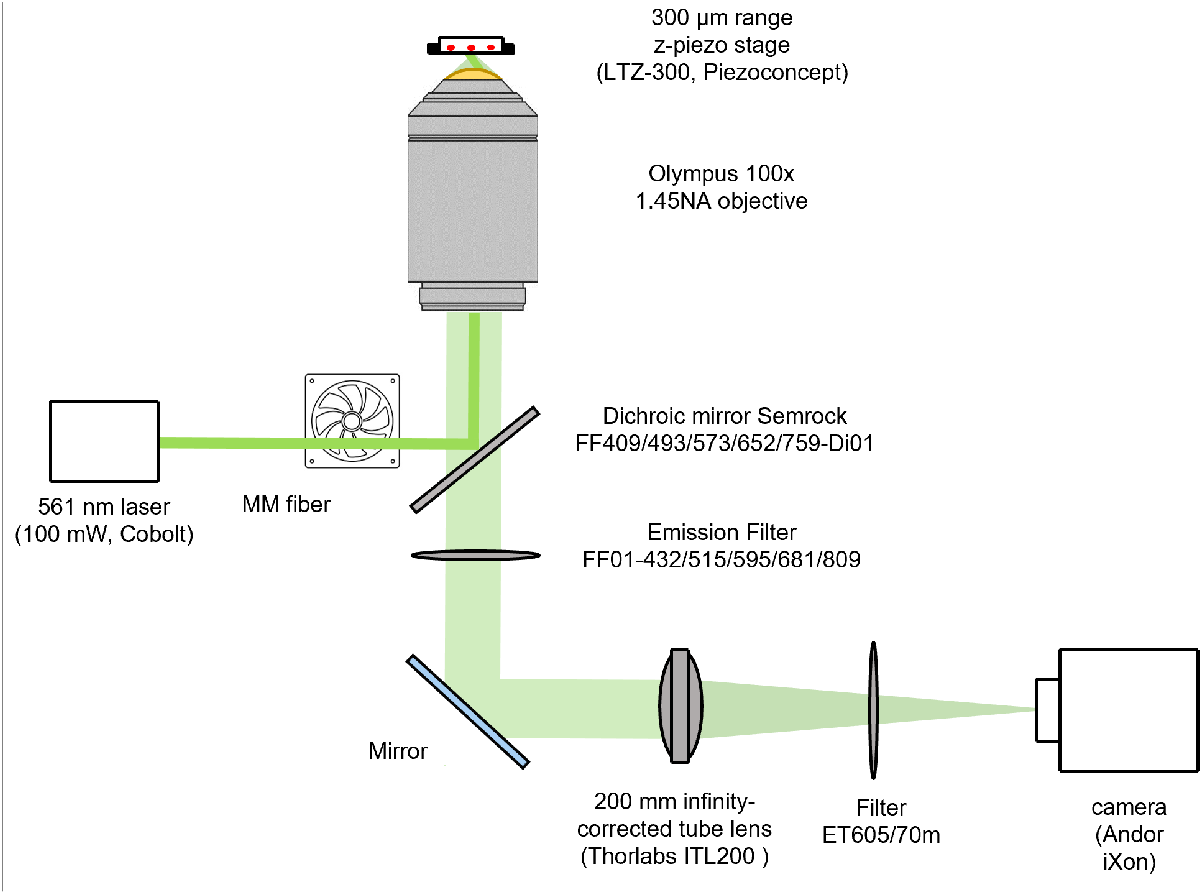
Optical setup for single color experiment with CF-568

### Sample Preparation and Immunofluorescence Staining

African green monkey kidney cells (COS-7) were cultured in DMEM-Glutamax (Gibco 10566016) supplemented with 10% FBS in a cell culture incubator (37°C and 5% CO2). Cells were plated at low confluency on ethanol-cleaned 25 mm #1.5 thickness round coverglass (VWR) for imaging. Prior to fixation, all solutions were pre-warmed to 37°C. 24h after plating, cells were pre-extracted for 30 s in 0.25% Triton X-100 (Sigma-Aldrich) in PHEM (60 mM PIPES, 25 mM HEPES, 10 mM EGTA, 4 mM MgSO4) washed in PHEM, fixed for 8 min in −20°C Methanol (Sigma-Aldrich), then washed 3 times with PBS. The samples were then blocked for 1h in 5% BSA, before being incubated for 1.5h at room temperature with 1:500 rat anti alpha-tubulin antibodies (abcam ab6160) in 1% BSA diluted in PBS-0.2% Triton (BSA-PBST), followed by 3 washes with PBST, and then incubated for 1h in BSA-PBST with a goat anti-rat CF-568 (Sigma SAb4600086) secondary antibodies, 1:500 For further imaging after the optimization was performed and for multicolor imaging, we used glutaraldehyde fixation based on the the protocol from (21), with a pre-extraction step of 30 seconds in PHEM-0.25% Triton + 0.1% glutaraldehyde, followed by fixation in PHEM-0.25% Triton + 0.5% glutaraldehyde for 8 minutes, and quenching in PBS-0.1% NABH4 for 8 minutes and finally 3 washes in PBS. The samples were then blocked for 1h in 5% BSA, before performing the immunostaining. We incubated the primary antibodies for 1.5h in BSA-PBST at room temperature followed by 3 washes with PBST and incubation with secondary antibodies for 1h in BSA-PBST. For single color imaging, we use two different primary antibodies: mouse anti alpha-tubulin (Sigma T6199) 1:500 and rat anti alpha-tubulin (abcam ab6160) 1:500, with the following secondary antibodies: (Fig 3,4) Goat anti-rat CF 568 (Sigma SAb4600086) 1:500, (Fig 5) Goat anti-rat Alexa Fluor 647 (Invitrogen A21247) 1:500 Horse anti-mouse Dylight 649 (Vectorlab DI-2649) 1:500, Goat anti-mouse CF 647 (Sigma SAb4600182) 1:500, (Fig 6) Goat anti-mouse CF 750 (Sigma SAb4600211) 1:250, Donkey anti-rat Dylight 755 (Invitrogen SA510031) 1:250, and Goat anti-rat CF 770 (Sigma SAb4600479) 1:250.

For 2-color imaging (Figure 7), we used Mouse anti alpha-tubulin (Sigma T6199) 1:500 and Rabbit anti clathrin heavy chain (abcam, ab21679) 1:500, and for secondaries Goat anti-mouse CF 647 (Sigma SAb4600182) 1:500 and Donkey anti-rabbit CF 568 (Sigma, SAB4600076) 1:500.

For 3-color imaging (Figure 8), we used Mouse anti alpha-tubulin (Sigma T6199) 1:500 and Rabbit anti clathrin heavy chain (abcam, ab21679) 1:500, and for secondaries goat anti-mouse CF 750 1:250 (Sigma SAb4600211) and Donkey anti-rabbit CF 568 1:500 (Sigma, SAB4600076). We finally added Alexa Fluor 647 Phalloidin (Thermofisher A22287) 1:200 one hour in PBS just before imaging, and rinsed twice in PBS.

The sample were imaged in an Attofluor imaging chamber (Invitrogen, A7816), with 1 ml of imaging buffer and another 25 mm round coverglass on top to limit air exchanges.

### Buffer preparation

- DABCO [1,4-di-azobicyclo-(2.2.2.)-octane] (D27802, Sigma) was dissolved in distilled water to make a 1 M stock solution, adding HCl 12M until all the powder was dissolved and the pH reached pH 8.0. The pH value is critical since it mostly determines the pH of the overall buffer. The stock solution was kept in the fridge in the dark for several weeks.
- DTT 1 M (43816, Sigma) was used directly as purchased and was kept in the fridge for several weeks.
- Sodium Sulfite (S0505, Sigma) was dissolved in PBS 10x to 1 M as in (15), and was kept at room temperature on the bench for several weeks.

We typically prepared 10 mL of buffer at a time, and we adjusted the pH using NaOH and HCl and a pH-meter (Mettler Toldeo FE20) and stored it in the fridge. The composition of all buffer conditions tested in this paper is given in Table S1.

### Data acquisition

For the buffer optimization of CF-568, we acquired 12,000 images with *≈* 50 ms integration time using the conventional amplifier of the EMCCD camera, did not add UV light (which limits the achievable density, but makes comparing conditions easier), and manually corrected z-drift. The camera and z-piezo were controlled using micro-manager (22). 561 nm laser power at the sample was typically between 1 and 2 kW.cm^*−*2^.

For further imaging with the optimized buffer, we acquired between 10.000 and 30.000 images using micro-manager, using active stabilization on the IX83 except at 750 nm since the dichroic used for focus stabilization prevents the transmission of the fluorescent signal. Laser power at 532 nm, 640 nm and 750 nm was typically between 1 and 4 kW.cm^*−*2^.

### Data Processing

Raw STORM image stacks were processed using using a FIJI macro that runs an analysis with Detection of Molecules (DoM) (23) and Thunderstorm (24), including drift-correction and grouping of consecutive localizations. Using a python script, the distributions of SNR and number of photons, as well as the density of fluorophores were averaged. To calculate this latter, the STORM image was divided in small bins of 30 nm*×* 30 nm, and the number of molecules in the image was divided by the total surface area of the bins occupied by at least one fluorophore. (The two scripts are available at (25)). FRC (26) values shown in figure 3 and 4 were calculated using the BIOP FIJI plugin (27) with DoM localizations exported in Thunderstorm format to create two images using the odd and even localizations modified from (28, 29), which is also available at (25). The diffraction limited images shown in figure 3 and 4 are the standard deviation of the raw STORM data, computed using FIJI.

## Results

### Optimization of CF-568 blinking

We first started with a buffer composition of 30 mM sulfite and 30 mM DTT, which is close to what was demonstrated to work with Alexa-647 with MEA instead of DTT (15), and looked at the influence of both pH and DABCO concentration. The mean SNR, photon count and density of molecules per dataset were studied using 11 buffers within a pH ranges of 7-9 and and a concentration range of 0-150 mM for DABCO (figure 2-a1,b1 & c1). A maximal point was reached for the three parameters for the buffer with a pH value 7.7 and DABCO concentration 65 mM. This pH value of 7.7 agrees with the optimal value of 7.5 found for STORM imaging of Alexa-647 when using DTT as a reducing agent (18).

**Fig. 2.**
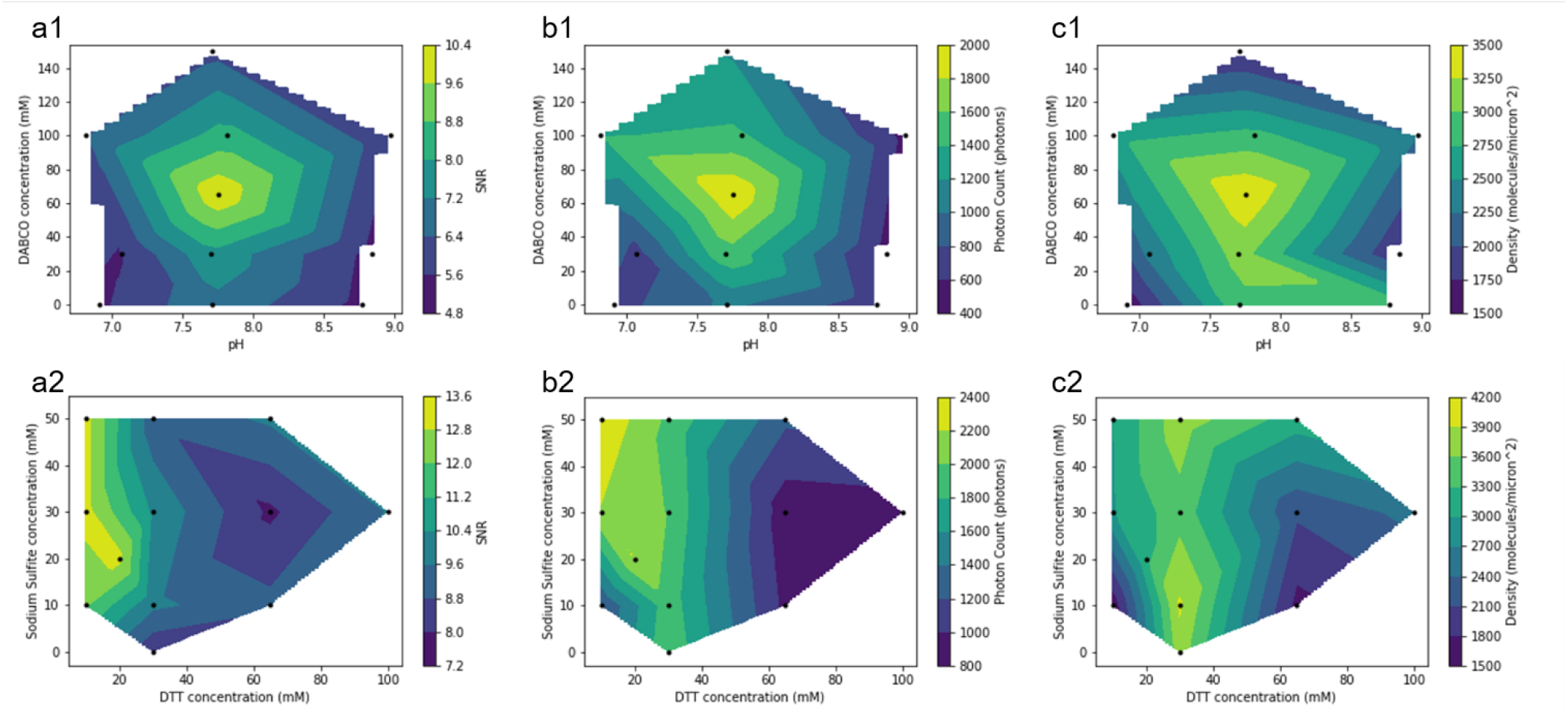
SNR, density and photon count of CF-568 as a function of the buffer composition for the 22 buffers tested for the optimization. (a1) SNR (b1) Photon counts and (c1) molecular density for 11 buffers with Sulfite concentration of 30 mM, DTT concentration of 30 mM, pH value given on the x-axis, and DABCO concentration on the y-axis. (a2) SNR (b2) Photon counts and (c2) molecular density for 12 buffers with DABCO concentration of 30 mM, a pH of 7.7, DTT concentration given on the x-axis, and Sulfite concentration on the y-axis.

We then fixed the pH to a value around 7.7 and the DABCO concentration to 65 mM and studied the influence of DTT and sodium sulfite concentrations by preparing 12 buffer with a concentration range of 10-100 mM for DTT and 0-50 mM for Sodium Sulfite (figure 2-a2,b2 & c2). The photon count and the density were highest for buffers with DTT concentration between 20 and 30 mM except when the sodium sulfite concentration was less than 10 mM. The buffers with 10 mM of DTT and 30 or 50 mM Sulfite also showed high SNR values but were ignored because of the high level of discontinuities observed in the final images despite their good brightness. Hence, the optimal range was found for buffers with a DTT concentration around 20-30 mM and having a sodium sulfite concentration above 10 mM.

To make sure we indeed reached an optimum for the chemical conditions of our buffer, we changed again the pH and DABCO concentration for a fixed DTT and sodium sulfite concentrations of 20 mM. We first fixed the pH to 8, and prepared 4 buffers with DABCO concentration in the range 0-100 mM. The same experiment was done using a DTT concentration of 30 mM and a sodium sulfite concentration of 10mM. In both experiments, the optimal concentration of DABCO was found to be 65 mM. In a similar manner, the concentrations of DTT and sodium sulfite were fixed to 20 mM and the DABCO concentration was fixed to 65 mM. Testing 3 buffers having respective pH values around 7, 8 and 9 clearly showed better results for the buffer with pH 8. We concluded that DTT and sodium sulfite concentrations around 30 mM, a DABCO concentration around 65 mM, and a pH value around 8 are optimal for CF-568 blinking.

We then tested this protocol on another microscope equipped with an active stabilization system and a single mode laser excitation at 532 nm (see Methods). We once again imaged microtubules, and obtained convincing results. As can be seen in figure 3 (a), the raw images show bright isolated molecules, and imaging can be performed for a long enough time to achieve a high density of molecules, resulting in a STORM image (figure 3 c) with a much better resolution than the diffraction-limited image reconstructed from the same dataset (figure 3 b). To estimate the resolution of our image, we performed a FRC (26) calculation which yielded a resolution of 11.5 nm. Convinced that our imaging protocol worked well enough to do high quality STORM imaging, we decided to test how stable our buffer was.

**Fig. 3.**
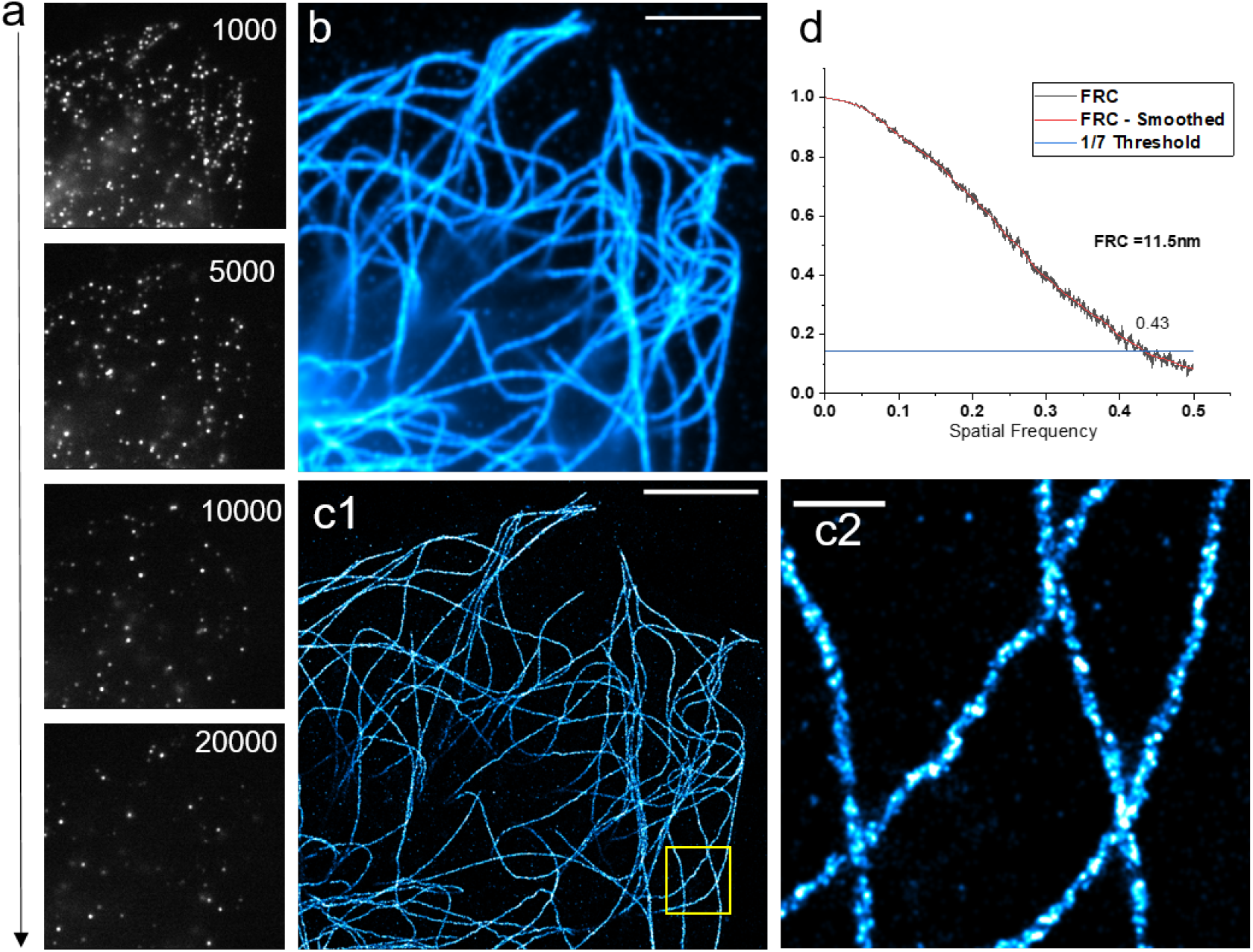
*α*-tubulin stained with CF-568 in our optimized buffer. (a) Raw camera frames with frame number indicated in the top-right corner(b) diffraction limited image (c) STORM image, reconstructed from ≈ 1.4 million molecules localizations in 26,000 frames. Scalebar: 5*μ*m, 500 nm in inset. (d) FRC curve for the dataset, giving an FRC resolution of 11.5nm

### Testing the buffer stability

We froze 10 mL of our optimized buffer, and left it at −20°C for a week. We then left it overnight in the fridge to thaw, and performed imaging the next day, as well as 4 days later (keeping the buffer in the fridge once thawed). We noticed no strong changes in the performance of the buffer (See fig 4) enabling the preparation of a large batch of buffer followed by aliquoting and freezing, guarantying reproducible performances throughout a STORM project.

**Fig. 4.**
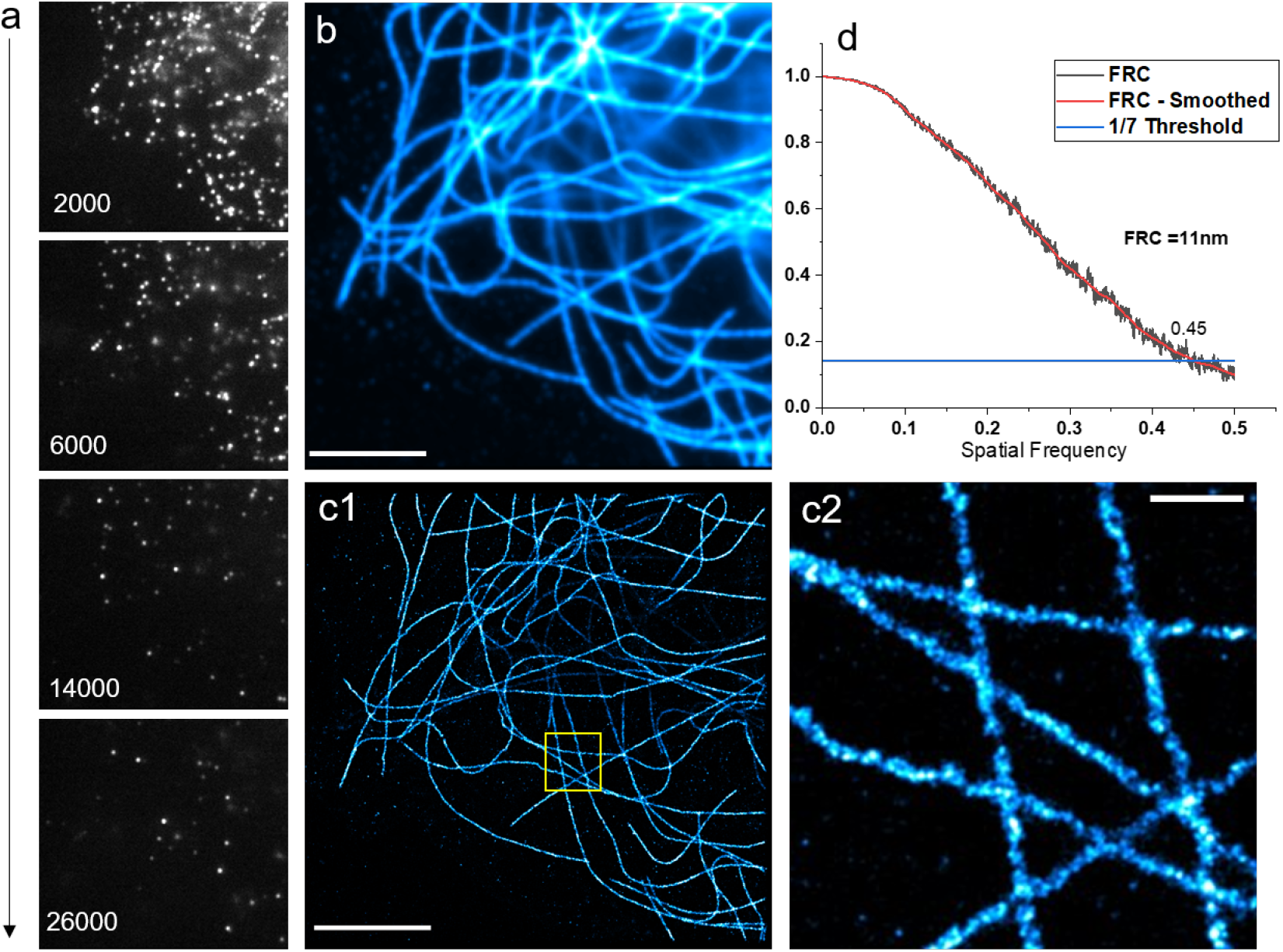
*α*-tubulin stained wit CF-568 in our optimized buffer after freezing 1 week and thawing overnight in the fridge and waiting 4 days (a) Raw camera frames with frame number indicated in the bottom-right corner(b) diffraction limited image (c) STORM image, reconstructed from ≈ 1.1 million molecules localizations in 30000 frames. Scalebar: 5*μ*m, 500 nm in inset. (d) FRC curve for the dataset, giving an FRC resolution of 11nm

### Testing 640 nm excited fluorophores in the optimized Buffer

Our initial assumption was that the ingredients of the buffer were chosen to insure good blinking of Alexa-647, but we had to confirm this. We therefore immunostained *α*-tubulin once again, and tested 3 popular far-red dyes: Alexa-647, Dylight-649 and CF-647, and obtained high quality images for all three, as shown in figure 5.

**Fig. 5.**
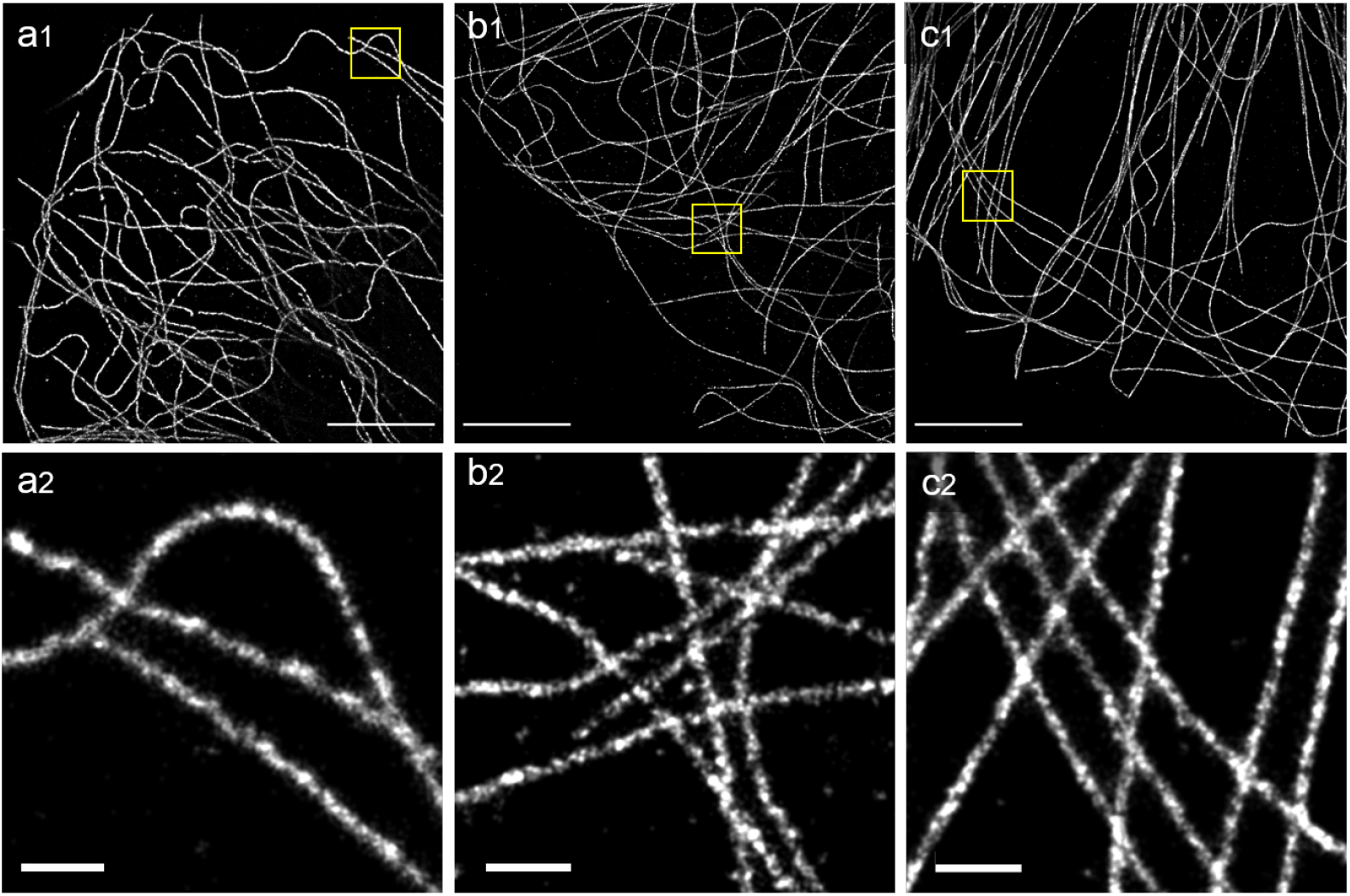
Test of several 640nm-excited fluorophores on *α*-tubulin. (a) Alexa-647 (b) Dylight-649 and (c) CF-647. Scalebars: 5*μ*m, 500 nm in inset

We quantified the average SNR, Photon counts and density of molecules for all three dyes in our buffer, and the value are given in Table 1. Whiles the differences in SNR are quite large (Possibly due to different amounts of background in the different images) all three dyes display high photon number, as well as high molecular density resulting in high quality images. We did notice however that in this wavelength range, two populations of molecules seemed to co-exist, as we could see on the raw data some bright short-lived molecules as well as some dim longer-lived ones.

**Fig. 6.**
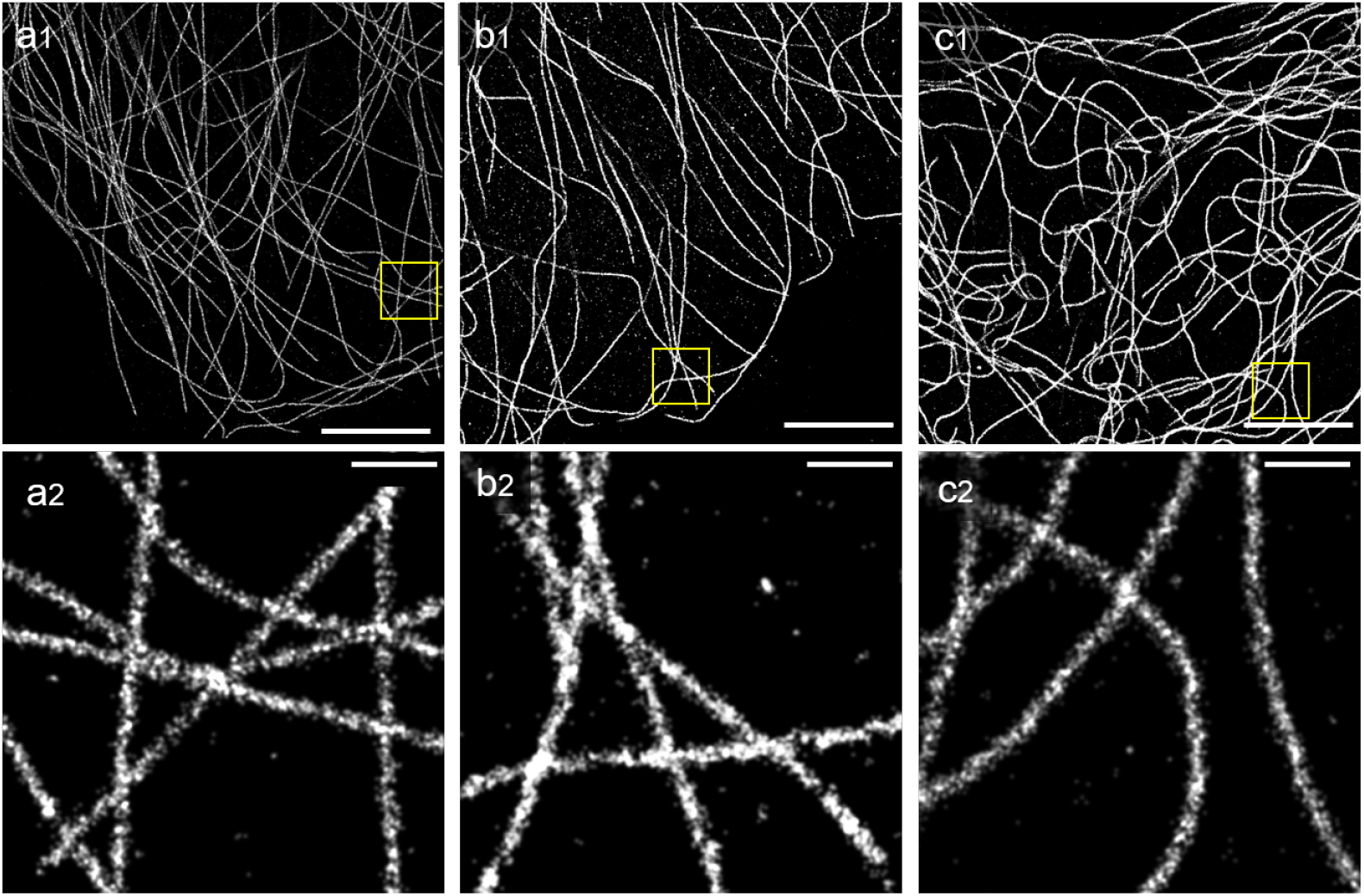
Test of several 750 nm-excited fluorophores on *α*-tubulin. (a) CF-750 (b) Dylight-755 and (c) CF-770. Scalebars: 5*μ*m, 500 nm in inset

**Table 1.** Comparison between the three different red dyes tested. The density is in molecules per *μm*_2_ excluding empty pixels. (see Methods)

| Dye | SNR | Photon count | Density |
| --- | --- | --- | --- |
| Alexa-647 | 9.8 | 9087 | 7134 |
| CF-647 | 18.9 | 5976 | 5716 |
| Dylight-649 | 13.9 | 7640 | 5091 |

### Testing 750 nm excited fluorophores in the optimized Buffer

Having made sure the usual 640 nm excited fluorophores worked, we tested several 750 nm excited dyes. These dyes have generally been shown to blink well with 640 nm excited fluorophores (7), but provide fairly low photon counts. We tested 3 dyes: Dylight 755, CF-750 and CF-770. In all 3 cases, we could reconstruct good quality images, as can be seen in figure 6 but the photon counts and SNR were as expected lower than for the other 2 colors (see Table 2).

**Table 2.** Comparison between the three different far-red dyes tested. The density is in molecules per *μm*_2_ excluding empty pixels

| Dye | SNR | Photon count | Density |
| --- | --- | --- | --- |
| CF-750 | 9.3 | 1647 | 2402 |
| Dylight-755 | 9.2 | 2900 | 2470 |
| CF-770 | 7.3 | 2100 | 2620 |

### Multicolor Imaging

Having just demonstrated that the far-red fluorophores tested worked well in our buffer, we decided to perform some multicolor STORM imaging.

#### 2-color Imaging

The most common laser combination on STORM microscopes is 532 nm and 640 nm, so we first tested how well our buffer behaved for a typical 2-color experiment, and chose the popular combination of microtubules (*α*-tubulin) and clathrin (30). We stained microtubules with CF-647, and Clathrin with CF-568, and imaged the two dyes sequentially starting with CF-647. Figure 7 shows images comparable to that obtained with a single fluorophore, demonstrating negligible cross talk.

**Fig. 7.**
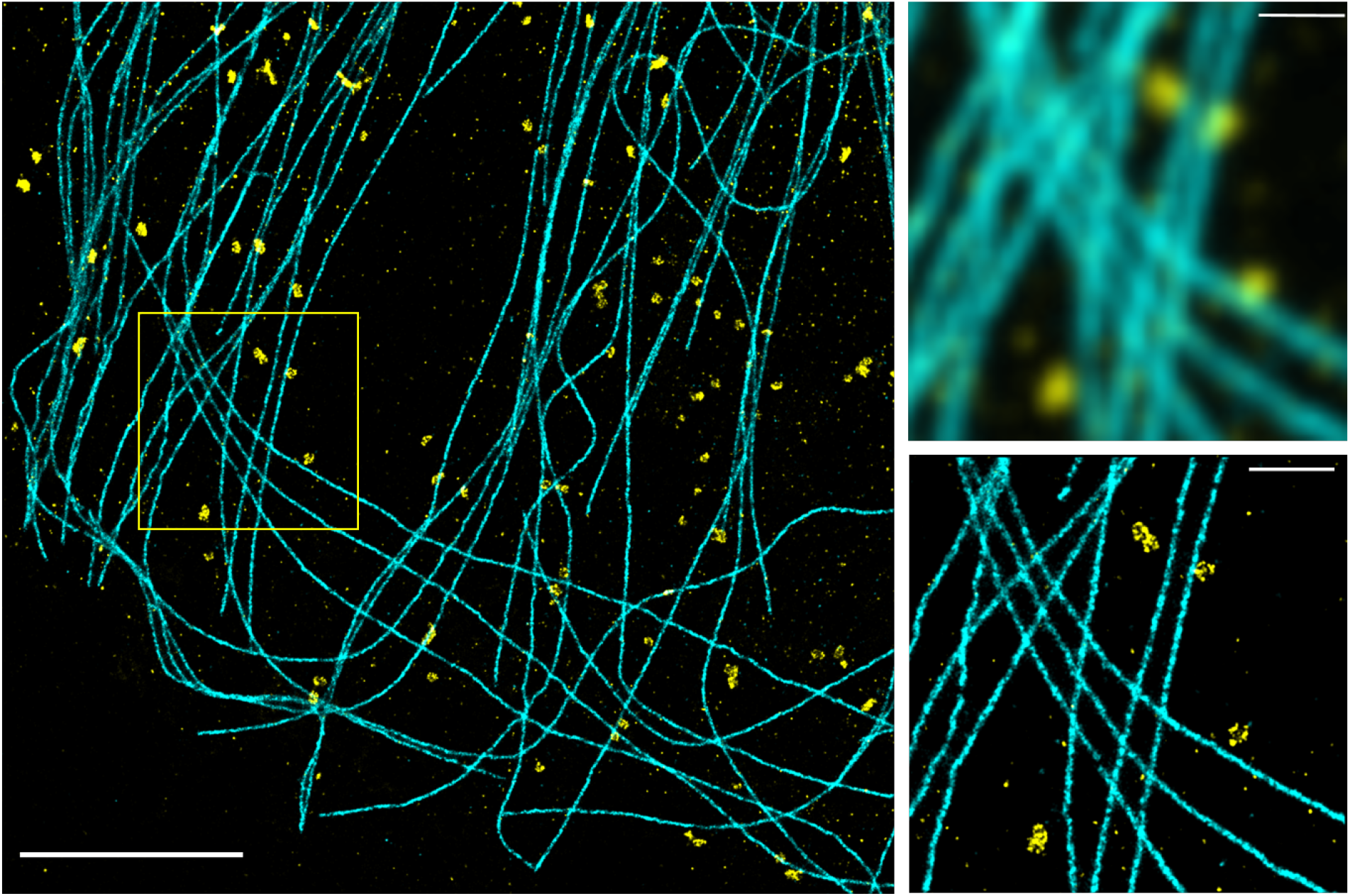
2-color STORM imaging of microtubules and Clathrin stained with CF-647 an CF-568 respectively. Scalebar: 5*μm*. Inset: approximated diffraction limited image and STORM image of the region indicated in the yellow box. Scalebar: 1*μm*

#### 3-color Imaging

Finally, we decided to test 3-color imaging, so we stained the cytoskeleton of our COS-7 cells: *α*-tubulin with CF-750, Clathrin with CF-568 and Actin with Alexa-647 (see Methods). We once again performed sequential imaging, starting with the reddest fluorophore. Figure 8 shows that this combination works well for multicolor imaging, with minimum cross-talk, good localization precision and good density for all three fluorophores. We also tested a few more fluorophores, some of which performed well enough to reconstruct a STORM image (see Table S2 and Figure S1). In particular, Alexa-532 and CF-680 worked in our buffer, which means spectral un-mixing (8, 11, 12) should be an option to increase the number of colors to 5.

**Fig. 8.**
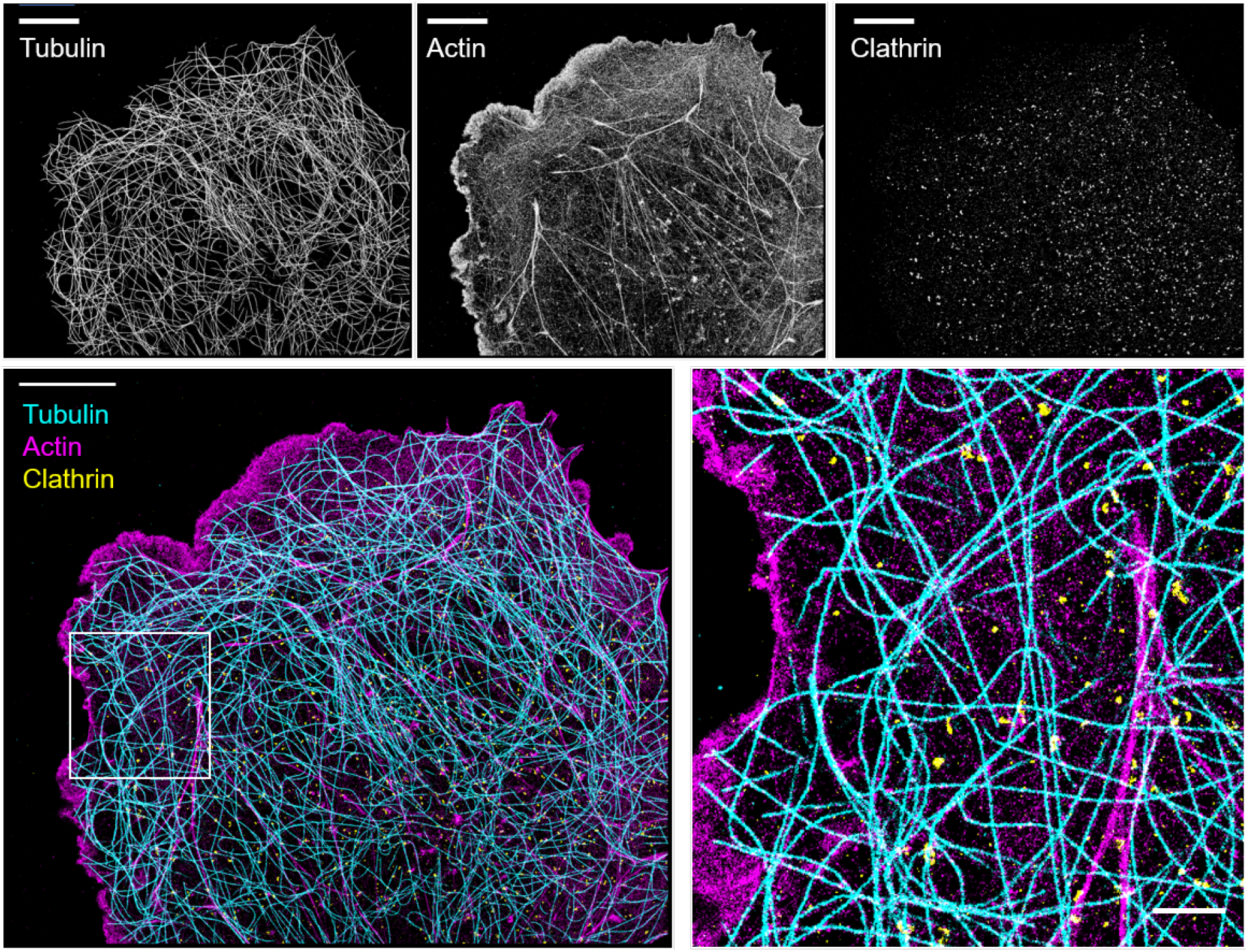
3-color STORM imaging.Top: From left to right: *α*-Tubulin (CF-750), Actin (Alexa-647), & Clathrin (CF-568). Bottom: composite image, with *α*Tubulin in cyan, actin in magenta and Clathrin in yellow. Scalebar=10*μ*m, 2*μ*m in inset

## Conclusions

We have shown that our optimized buffer composition allows 3-color STORM imaging using multiple fluorophore combinations, and in particular that it enables high quality 2-color imaging using Alexa-647 and CF-568, using both 561 nm and 532 nm excitation. This buffer is stable for more than a week, can be frozen and stored at −20°C, and is relatively inexpensive to make (see Supplementary Note). We do not claim that the buffer we presented is better than other buffers, and for 2-color imaging for example the combination of Alexa-647 (or Dylight-649, or CF-647) and CF-750 with an optimized buffer (9, 10) is probably preferable. However, a lot of microscopes are not equipped with a 750 nm laser line, in which case our protocol provides good performances for both Alexa-647 and CF-568. Our protocol also appears particularly well-suited for STORM imaging of large sample with light-sheet illumination that requires large amounts of buffers. Maybe more importantly, we believe that this stable buffer provides a very good starting point for further optimization, for example by replacing/adding another reducing agent like MEA (31), or TCEP (9, 32), adding other chemicals such as Propyl Gallate (16, 17) Ascorbic Acid (5), or Trollox (33). Indeed, as a proof-of-principle experiment, we tested adding COT (34) to our buffer with CF-750, and saw an increase in the brightness of the molecule consistent with previous reports (See Figure S2).

A similar optimization was recently used to improve low-power single-color STORM with Alexa-647 (18), so we expect a broad family of buffers optimized for specific situations to be developed. This approach, in conjunction with the development of new fluorescent dyes optimized for STORM (35) should lead to further improvements for multicolor imaging.

### Limitations

- We performed our optimization with a simple microscope using a simple sample preparation, which increases the experimental variability. As we measured a fairly continuous dependence of image quality as a function of the different concentrations this did not prevent us from performing the optimization but more robust sample prep and stabilized microscopes would certainly help.
- We also kept the image analysis very simple to see if such a brightness/density based-approach was sufficient, but or a fully-automated image-based optimization, more advanced analysis would be required to avoid artifacts.
- We determined the recipe for our buffer using a fairly simple optimization scheme, with only 4 parameters (concentration of 3 chemicals plus pH) on a single fluorophore, so we expect that further improvements are possible. For example repeating our measurements on 640 nm and 750 nm-excited fluorophores might yield a better 3-color buffer. Indeed, we noticed in our 640 nm excited mages the appearance of two populations of molecules, bright ones and dim ones which might be an issue for some sensitive measurements, but might be resolved by tweaking the concentrations of the chemicals.
- We tried to limit the use of toxic chemicals, but further work is needed to obtain a truly safe imaging buffer.
- Finally we did not investigate the photophysical mechanisms behind the blinking we observed (36, 37). Understanding these mechanisms should allow us to further optimize the buffer composition in a more rational manner.

## Data availability

- The raw STORM data for the main text and supplemental data are available on Zenodo (38)
- The FIJI macro used to convert the raw data into STORM images in on github (25), along with the Python code used to quantify photon counts, SNR and density.

## Author Contributions

CF-568 buffer optimization: V.A. prepared the samples, performed STORM imaging and analyzed the data. H.B. performed preliminary experiments and helped set the parameters. Multicolor imaging: N.O. prepared the samples, V.A. and N.O performed STORM imaging and V.A. analyzed the data. V.A and N.O. wrote the article. N.O. supervised the project.

## ACKNOWLEDGEMENTS

N.O. acknowledges funding from CNRS (Tremplin@INP 2019 and Tremplin@INP 2021). We thank Sandrine Leveque-Fort for the COS-7 cells and Maxime Mauviel for his help with cell culture, abbelight for the the gift of Clathrin secondary antibody, Nicolas David for Alexa-647 Phalloidin, Beatrice Durel for the Alexa-532 and Cy3 antibodies, Vectorlab for the gift of the Dylight-649 antibody, Biotium for the gift of CF-532, CF-660C and CF-770 secondary antibodies. We thank Debora Keller-Olivier and Dorian Noury for critical reading, and all the members of the “nano” group at LOB for scientific and technical discussions. We thank Ricardo Henriques for the Latex template.

## Notes

### Competing Interest Statement

The authors have declared no competing interest.

### Summary of Updates

- Modified Figure 6 - Added a link to the raw data - added a table with all the buffers tested - fixed a few typos

