## Supplementary data for "An Optimized Buffer for Repeatable Multicolor STORM"

### An Optimized Buffer for Repeatable Multicolor STORM
### Supplementary Information

Vaky Abdelsayed, Hadjer Boukhatem & Nicolas Olivier

#### 1 List of all buffer conditions tested

| name | pH | DABCO | Sodium Sulfite | DTT |
|---|---|---|---|---|
| b1 | 6.9(M) | 0 | 30 | 30 |
| b2 | 7.1(M) | 30 | 30 | 30 |
| b3 | 6.8(M) | 100 | 30 | 30 |
| b4 | 7.7(M) | 0 | 30 | 30 |
| b5 | 7.7(M) | 30 | 30 | 30 |
| b6 | 7.7(M) | 65 | 30 | 30 |
| b7 | 7.8(M) | 100 | 30 | 30 |
| b8 | 7.7(M) | 150 | 30 | 30 |
| b9 | 8.8(M) | 0 | 30 | 30 |
| b10 | 8.8(M) | 30 | 30 | 30 |
| b11 | 9.0(M) | 100 | 30 | 30 |
| b12 | 7.7(E) | 65 | 0 | 30 |
| b13 | 7.7(E) | 65 | 10 | 10 |
| b14 | 7.7(E) | 65 | 10 | 30 |
| b15 | 7.7(E) | 65 | 10 | 65 |
| b16 | 7.7(E) | 65 | 20 | 20 |
| b17 | 7.7(E) | 65 | 30 | 10 |
| b18 | 7.7(E) | 65 | 30 | 65 |
| b19 | 7.7(E) | 65 | 30 | 100 |
| b20 | 7.7(E) | 65 | 50 | 10 |
| b21 | 7.7(E) | 65 | 50 | 30 |
| b22 | 7.7(E) | 65 | 50 | 65 |
| b23 | 7.7(E) | 0 | 20 | 20 |
| b24 | 7.7(E) | 0 | 10 | 30 |
| b25 | 7.7(E) | 30 | 20 | 20 |
| b26 | 7.7(E) | 30 | 10 | 30 |
| b27 | 7.8(E) | 100 | 20 | 20 |
| b28 | 7.8(E) | 100 | 10 | 30 |

Table S1: Composition of the buffers tested in figure 2.
Concentrations are given in mM. (M): measured - (E): Estimated.
Buffer b6 (highlighted in cyan) is the one used in the rest of the study

#### 2 Test with additional fluorophores

We tested the buffer with some additional dyes, some of which had already been tested in other buffers such as iFLuor647 [1] to get a better overview of the possible multicolor combinations. Our tests are summarized in table S2 and some representative images of the dyes deemed compatible are shown in Figure S1.

| Dye | Reference | Blinking | Misc. |
|---|---|---|---|
| Alexa-532 | Invitrogen A-11002 | ++ | see fig |
| CF-532 | Biotium 20365 | ++ | see fig |
| Alexa-555 | Invitrogen A-31570 | - | |
| Cy3 | Invitrogen A-10522 | - | |
| Dylight 550 | Invitrogen SA5-10039 | - | |
| iFLuor647 | Abcam ab176759 | + | see fig |
| CF-660C | Biotium 20050 | + | see fig |
| CF-680 | Biotium 20065 | ++ | see fig |
| IRDye680RD | Odyssey 926-68071 | + | see fig |

Table S2: Other dyes tested in the optimized buffer

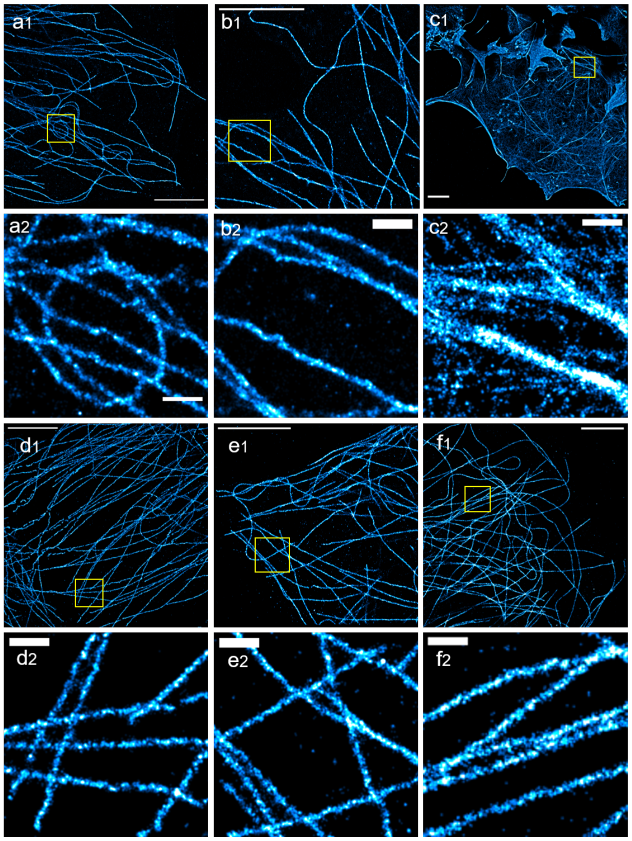


Figure S1: Other fluorophores tested that can be used for STORM imaging. (a) CF-532 (b) Alexa-532 (c) iFluor647 (d) CF-660 (e) CF-680 (f) IRDye680RD. All tests performed on $\alpha$-tubulin (mouse or rat primary, see methods) except iFluor647-phalloidin. Scalebars: $5\mu$ m, 500 nm in inset

### 3 Test with added COT

We added 2 mM COT (Sigma, 138924, dissolved to 200 mM in DMSO) to our buffer, and observed an enhanced number of photons for the dye CF-750, consistently with previous reports [2, 3, 4].

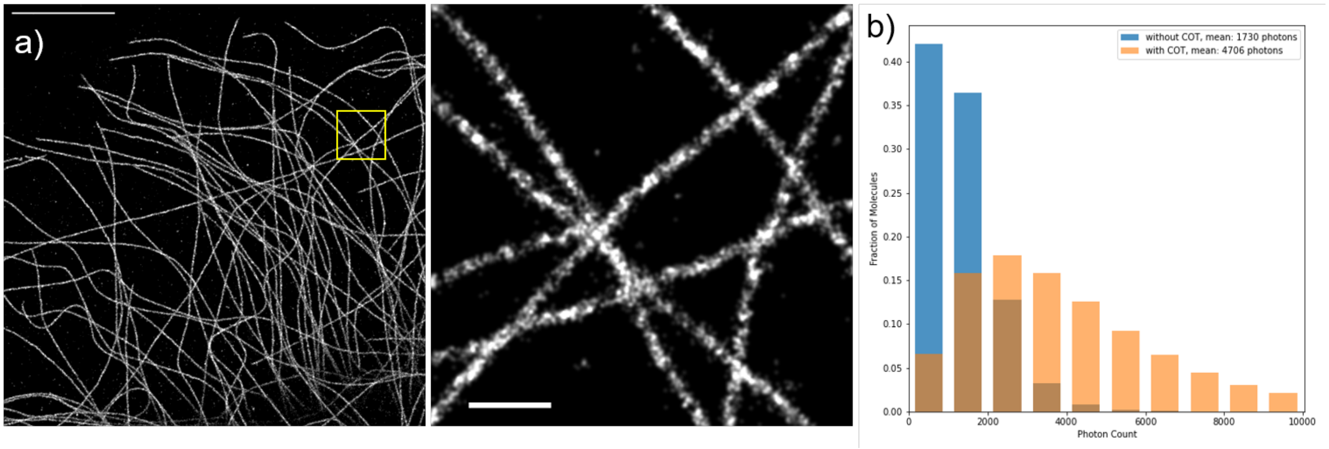


Figure S2: CF-750 with our optimized buffer complemented with 2 mM COT. (a) STORM reconstruction scalebars = 5$\mu$m (left) and 500 nm (Inset, right) and (b) Histograms for the photon distribution with and without adding COT

### 4 Cost of an Experiment

We use 1 ml of buffer per experiment. We estimate the cost (From Sigma-Aldricht, in France) as follow:
- We purchased DTT as 10 mL solution at 1 M for ≈ 25€, so enough for 330 mL of buffer, so less than 8 cents/mL. Buying it in larger quantities could easily half the price.
- We purchased Sodium Sulfite as 250 g of powder (mW:126, so 2M) for ≈ 25€, enough for more than 60L at 30 mM, or a cost of less than a cent per experiment
- We purchased DABCO as 25 g (mW:112.17 g/mol, so 0.223 M) for ≈ 25€, so enough for 3.4 L at 65 mM, or a cost of less than a cent per experiment

So overall, the cost is below 10 cents per experiment, and can be further reduced by purchasing larger quantities of reagents or using lower amounts of buffer.
